# Seabird traits and seasonality modulate nutrient dynamics of terrestrial and marine habitats on atolls

**DOI:** 10.1101/2023.12.03.569819

**Authors:** Jennifer Appoo, Nancy Bunbury, Jake Letori, Aurelie Hector, Annie Gendron, Nicholas A.J. Graham, Gerard Rocamora, Matthieu Le Corre, Sébastien Jaquemet

**Affiliations:** UMR ENTROPIE, Université de La Réunion, 97744 Saint Denis Cedex 9, La Réunion, France; Seychelles Islands Foundation, Victoria, Mahé, Seychelles; Centre for Ecology and Conservation, University of Exeter, Cornwall Campus, Penryn, UK; Island Conservation Society, Pointe Larue, Mahé, Seychelles; Lancaster Environment Centre, Lancaster University, Lancaster, UK; Island Biodiversity and Conservation Centre, University of Seychelles, Anse Royale, Mahé, Seychelles

**Keywords:** Island ecosystems, Marine subsidies, Seagrass, Seychelles, Sooty terns, Red-footed boobies, Stable isotopes, Tropical seabirds

## Abstract

Marine nutrients underpin productivity and functioning of oceanic island ecosystems. Seabirds represent a primary source of marine nutrients. In tropical regions, some of the largest seabird populations nest on atolls, yet there is limited information available on seabird contributions to atoll ecosystem nutrient dynamics. To investigate the spatial and seasonal dynamics of seabird contributions, we assessed seabird colonies of different taxa, including red-footed boobies and terns, nesting on separate islands of Farquhar Atoll, Seychelles. We assessed nutrient concentrations of guano, soil, coastal plants and nearby seagrass in seabird colonies and at a control island with no seabirds, during the wet and dry seasons. Sooty terns contributed the highest quantities of nutrients, estimated at 71.2 N tonne.yr^-1^ and 52.2 P tonne.yr^-1^. Seabird-derived nutrient transfer occurred year-round from seabird colonies to soil, coastal plants and seagrass. Soil macro- and inorganic nutrients were higher in the high-density, ground-nesting tern colony and during the dry season, coinciding with the breeding period of sooty terns. Both red-footed booby and tern colonies maintained high nitrogen levels (%N) in coastal plants year-round, while phosphorus levels (%P) did not differ between islands or seasons. Seabird-derived nitrogen reversed nitrogen limitation (log C:N) of seagrass during the dry season. We provide the first insights into seabird nutrient contributions to atoll ecosystems in Seychelles, with recommendations for seabird conservation to boost and support atoll and island ecosystem resilience. Our results from a relatively undisturbed atoll serve as a baseline with which more impacted atolls and future changes can be assessed.

## INTRODUCTION

Resource subsidies are critical in shaping island biodiversity and functioning (Polis et al., 1997; Obrist et al., 2020). Isolated from continental landmasses, oceanic islands receive organisms, detritus and nutrients mostly from the surrounding marine environment (Polis & Hurd, 1996). Islands are prime nesting and roosting habitats for seabirds, making seabirds a principal source of marine subsidies (Smith et al., 2011). Seabirds act as bio-vectors, transferring and depositing marine nutrients, mainly in the form of guano, from oceanic feeding habitats to their island breeding colonies (Otero et al., 2018). In tropical regions, some of the largest breeding seabird colonies occur on atolls (Berr et al., 2023). On atolls, seabird-derived nutrients sustain terrestrial plant communities (Schmidt et al., 2010; Young et al., 2010) and, when transferred to nearshore reefs via run-off, enhance biomass and growth of corals and fish (Graham et al., 2018; Benkwitt et al., 2019; Savage, 2019). These links have highlighted the major role seabird subsidies play in atoll biodiversity (Lorrain et al., 2017; Benkwitt et al., 2020) and ecosystem services (Plazas-Jiménez & Cianciaruso, 2020), and hence the need to better understand seabird contributions to atoll terrestrial and marine ecosystems (Berr et al., 2023). Despite this importance, studies of seabird effects on atolls remain limited and are often underrepresented in global meta-analyses (Grant et al., 2022; Van Der Vegt & Bokhorst, 2023).

Seabirds are taxonomically diverse, with substantial variation in their spatial and temporal nutrient subsidies (Pascoe et al., 2022). The dynamics of seabird subsidies are therefore governed by species-specific traits related to foraging and breeding. For example, nutrient content of guano depends on diet, nutrient distribution is determined by nesting location and density, and nutrient quantity is influenced by biomass and breeding duration (Smith et al., 2011). Many seabirds are also seasonal breeders, which dictates timing of nutrient deposition. Ultimately, these traits influence ecosystem responses to seabird subsidies, and although these relationships have been explored for temperate or polar seabirds (Zwolicki et al., 2013, 2016), they are relatively unexplored for tropical species. The behaviour of tropical seabirds differs markedly from seabirds in temperate or polar regions (Weimerskirch, 2007). For example, tropical seabirds have broader diets and feeding strategies (Catry et al., 2009), and they often have non-seasonal and protracted breeding cycles (Catry et al., 2013; Carr et al., 2021). Investigations of seabird subsidies on atolls have mainly compared seabird effects between high or low seabird densities, regardless of taxonomy (McCauley et al., 2012; Benkwitt et al., 2021). How species-specific seabird foraging and breeding traits influence tropical seabird nutrient provisioning, and the broader implications for atoll nutrient dynamics, however, has not yet been studied.

Here, we report on a quantitative and qualitative assessment of seabird-derived nutrients to atoll terrestrial and marine ecosystems on the relatively undisturbed Farquhar Atoll in Southern Seychelles. We asked the following questions: (1) What is the nutrient input to atolls from seabird colonies of different taxa? (2) Are seabird-derived nutrients transferred to terrestrial and nearshore environments and how does this vary across taxa and seasons? (3) How do nutrient levels in terrestrial and nearshore habitats differ between seabird colonies and seasons? And (4) Do seabird-derived nutrients reverse nutrient limitations and increase growth rates of coastal and marine plants? To address these questions we sampled guano, soil and coastal plants, representing the main medium and basal components of nutrient transfer from seabird guano to atoll communities, respectively. To assess nutrient provisioning from seabird colonies to the nearshore environment we sampled seagrass, a prominent marine macrophyte of atolls. Seagrasses incorporate nutrients from the surrounding waters over a relatively long time frame, making them ideal indicators of nutrient availability in nearshore environments (Duarte, 1990; Allgeier et al., 2013). We discuss our results in the context of island seabird conservation.

## 2. METHODS

### 2.1. Study area

Farquhar Atoll (10°11′ S, 51°06′ E) lies in the Western Indian Ocean, ca. 770 km from Mahé, the main island of Seychelles, and 285 km northeast of Madagascar (Figure 1). Farquhar consists of 11 islands (total landmass 8 km^2^), the largest three being North Island, South Island and Goëlettes (see also Online Supplementary Material [OSM]). Anthropogenic activity on Farquhar is limited to a small settlement on North Island operating an ecotourism establishment, (10 beds, ca. 40 staff, Duvat et al., 2017). On North and South Islands, the beach vegetation consists of native coastal shrubs, which transitions into a mix of indigenous and introduced grasses and trees inland. In contrast, Goëlettes is treeless and covered in a low herb plant community, with coastal shrubs along the shores (Stoddart & Poore, 1970). There are large expanses of seagrass adjacent to the islands in the lagoon and reef flats, dominated by *Thalassodendron ciliatum* (Stokes et al., 2019).

**Figure 1.**
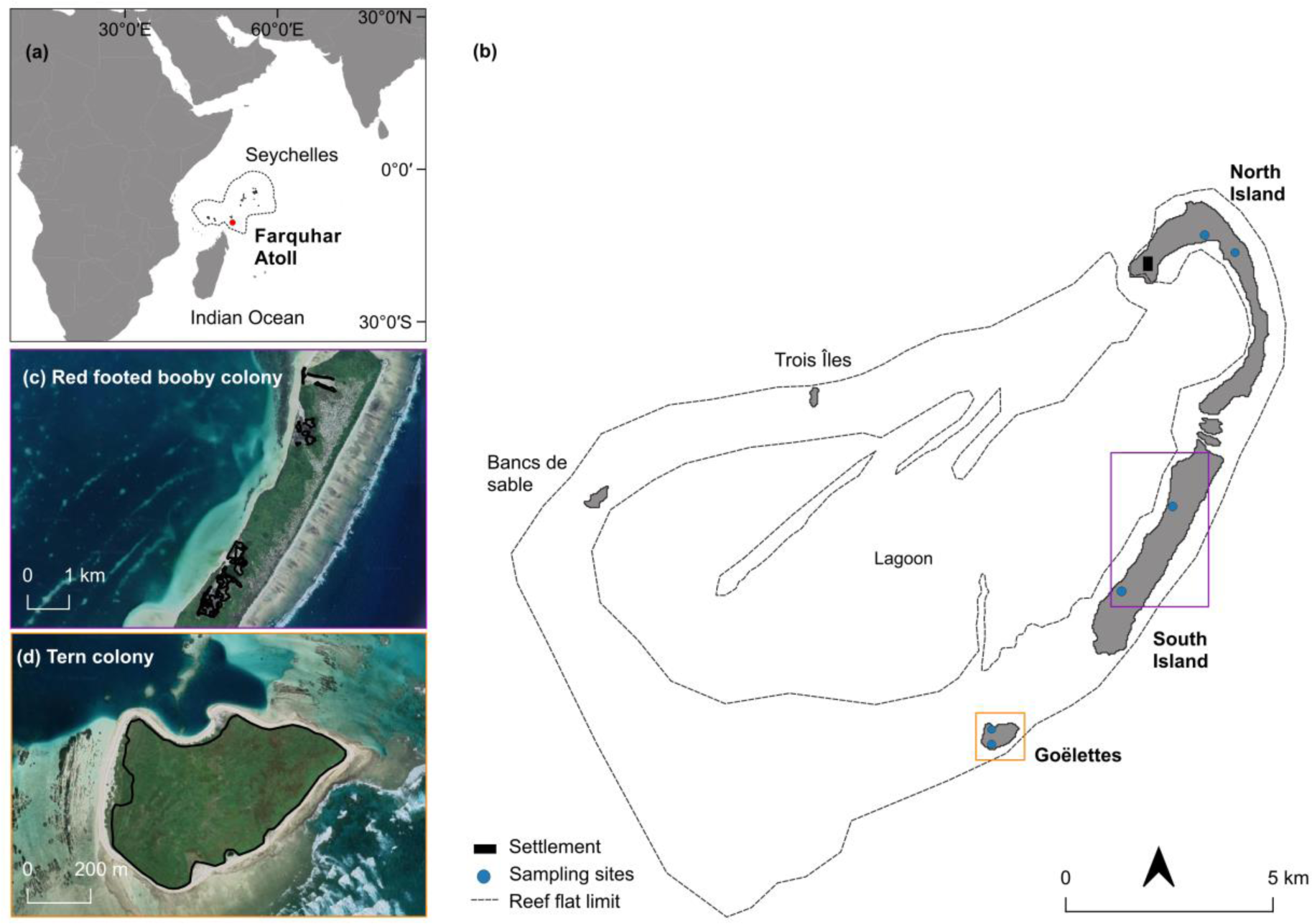
(a) Farquhar Atoll (in red), Seychelles, in the Western Indian Ocean; (b) Study islands and locations of sampling sites. Locations of (c) red-footed booby colony on South Island and (d) tern (comprising brown noddies and sooty terns) colony on Goëlettes, delimited in black lines.

Large breeding colonies of red-footed boobies *Sula sula*, brown noddies *Anous stolidous* and sooty terns *Onychoprion fuscatus*, are found on two separate islands (Duhec et al., 2017). Sooty terns and brown noddies nest on Goëlettes between May and October, and breed on the ground. Two red-footed booby colonies are located along the lagoon coastline and tidal inlets of South Island. Red-footed boobies breed year-round with peaks in March-April and November-December in nests around 1‒2 m from the ground. All three species prey mainly on pelagic fish and cephalopods (Weimerskirch et al., 2005; Catry et al., 2009). North Island, in contrast, has almost no breeding seabirds, attributed to the island’s historical use as the main centre for human settlement and coconut exploitation (Duhec et al., 2017). Most rain on Farquhar falls between November and April as a result of north-east monsoon winds. Between May and October, trade winds blowing from the south-east result in lower rainfall (Piggot, 1961).

### 2.2. Sampling design

We investigated the influence of seabird traits on nutrient dynamics using three separate islands and their breeding seabird species as a treatment group: (a) red-footed boobies on South Island, (b) terns, comprising brown noddies and sooty terns, on Goëlettes, and (c) North Island, with no breeding seabirds, as a control island. Because rainfall influences nutrient dissipation and availability in nearshore environments (Signa et al., 2021), we incorporated local seasonality by sampling in both the wet (March 2022) and dry seasons (August 2022). Full details of the sampling and analytical methods are given in the OSM.

### 2.3. Seabird nutrient input

To determine the quantity of nutrients delivered by seabirds, we collected fresh seabird droppings from the three breeding species within their colonies. Individual droppings were combined to obtain a minimum of 30 g wet weight per species for nutrient analyses and stored frozen until processing (Staunton Smith & Johnson, 1995). We determined macro- and micro-nutrient content of fresh guano for each species. Macronutrients consisted of total N, ammonium (N-NH_4_ ^+^), nitrate (N-N0 ^-^) and total P, while micronutrients comprised of iron (Fe), manganese (Mn), zinc (Zn) and copper (Cu). We estimated the total annual N and P input from seabird droppings using their concentrations measured in guano, excrement rates, population size and breeding metrics for each species, based on previously used methods (Riddick et al., 2018; Graham et al., 2018).

### 2.4. Terrestrial and marine sampling

We sampled soil and leaves of coastal vegetation in two habitat types on each island; coastal shrub habitat and low herbaceous habitat along transects parallel to the lagoon shore. On the two seabird islands, we sampled within the colonies. For the marine sampling, we collected leaves of seagrass adjacent to each island. We collected a sub-sample from each sample for isotope analysis and used the remaining sample for nutrient analyses. All samples for isotope analysis were stored frozen upon return from the field until further processing while the remaining soil, coastal plant and seagrass were air-dried. We sampled the same locations in both seasons.

### 2.5. Nutrient analyses

Nutrient levels and stoichiometric ratios provide useful insights into ecosystem functioning. For soil samples, we determined the pH, electrical conductivity (EC), total carbon (C), organic carbon (C org), N-NH_4_ ^+^ and N-NO_3_ ^-^, bioavailable phosphorus (P-a), cationic exchange capacity (CEC) and exchangeable cations (Ca^2+^, Mg^2+^, Na^+^ and K^+^). For coastal plants and seagrass, we assessed total C and organic C, respectively, and total N and P for both sample types. Levels of limiting nutrients (N and P) determine the quantity of nutrient subsidy and provide information about controls on ecosystem productivity (Polis et al., 1997). We derived stoichiometric ratios for coastal plants and seagrass as indicators of nutrient limitations (Sitters et al., 2015). In plants, reductions in nutrient limitations, determined by low C:nutrient ratios, are diagnostic of high plant nutrient quality (Sitters et al., 2015) and growth rate (Ågren, 2004, 2008). Nutrient analyses were conducted at the CIRAD-Reunion in the laboratory of agronomic analyses.

### 2.6. Isotopic analyses

Evidence on nutrient origin can be tracked through nitrogen isotopic ratios (δ^15^N) (Mizutani & Wada, 1988). Because they feed at high trophic levels in the marine environment, seabirds acquire a high δ^15^N signature, making δ^15^N a reliable tracer of seabird-derived nutrients (Pascoe et al., 2021). We determined δ^15^N on sub-samples of seabird guano, soil and plant material. Additionally, we determined carbon isotopic ratios (δ^13^C) on samples of coastal plants and seagrass. In plants, δ^13^C level is an indicator of physiological processes (Marshall et al., 2007), such as high photosynthetic activity or water stress (Mulder et al., 2011). Stable isotope analyses were conducted at Lancaster University, UK. Soil and seagrass samples were run twice, once after repeated acidifications with 1 M HCl to remove carbonates for δ^13^C analyses and once without this treatment for δ^15^N. Accuracy based on internal standards was within 0.2 ‰ *SD* for δ^15^N and 0.1 ‰ *SD* for δ^13^C.

### 2.7. Data analysis

All analyses were conducted in R version 4.3.0 (R Core Team, 2022). We tested for differences in baseline seabird-derived nutrients entering each colony by comparing guano δ^15^N between taxa. Shapiro-Wilk tests revealed non-normality in residuals, therefore we used Kruskal-Wallis test, followed by Dunn’s post-hoc tests with Bonferroni correction to identify the taxa. We assessed the transfer of seabird-derived nutrients using linear mixed models (LMMs) for soil, coastal plants and seagrass separately using *lme4* package (Bates et al., 2015). We tested for differences in δ^15^N as response variable, between both main and interaction effects of treatments (control, red-footed booby, tern) and seasons (wet, dry) as predictor variables. Sampling location was included as a random effect to account for repeated sampling. For soil and coastal plants, we pooled measurements of the two habitats since we were interested in the community-level response, and added habitat type as a covariate in the model. Where significant differences were detected, we performed post-hoc tests with Bonferroni adjustment to identify differing groups using the *emmeans* package (Lenth, 2023).

We used principal component analysis (PCA) to explore differences in soil properties and nutrient concentrations between treatment groups and season. PCA was conducted on scaled values and Euclidean distance matrix using the unconstrained redundancy analysis from the *vegan* package (Okanasen et al., 2018). We conducted permutational multivariate analysis of variance (PERMANOVA) based on Euclidean distance matrix to identify differences in soil parameters between treatments, seasons and their interaction. We then performed pairwise comparisons between each level of treatment group with the package *pairwiseAdonis* (Arbizu 2017), using 10,000 permutations and Bonferroni *p*-value adjustment.

To examine the influence of seabirds on foliar nutrient levels and ratios of coastal vegetation and seagrass, we ran separate LMMs for each nutrient type (N, P), ratio (C:N, C:P) and δ^13^C as response variables. Nutrient ratios were determined on a molar basis, and the log of ratios were used in all analyses and representation to avoid errors and misinterpretation (Isles, 2020). We tested for differences in predictor variables between both main and interaction effects of treatments and seasons, with sampling location as a random effect. Habitat type was included as a covariate for coastal plants, with pooled measurements from the two habitats. We performed post-hoc tests with Bonferroni adjustment between factor levels where significant differences were detected. Lastly, we plotted δ^15^N against %N to verify whether seabird subsidies (enrichment in δ^15^N) resulted in biologically relevant uptake of nitrogen %N in coastal plants and seagrass (Obrist et al., 2022).

For all LMMs, we computed the marginal and conditional R^2^ as goodness-of-fit metrics using the *MuMin* package (Barton, 2023). Marginal and conditional R^2^ describe the proportion of variance explained by the fixed effects only and fixed and random effects, respectively. For three of 13 models, sampling location (random effect) explained zero variance and led to a singular fit. Removing the random effect led to the same results so we kept it because of our repeated sampling. Model assumptions were verified using the protocol described by Zuur and Ieno (2016).

## 3. RESULTS

### 3.1. Nutrient input from seabird colonies of different taxa

Breeding sooty terns deposited the highest annual quantities of total N and total P to Farquhar Atoll, estimated at 71.2 tonne.yr^-1^ and 52.2 tonne.yr^-1^ (dry weight), respectively, followed by red-footed boobies (8.43 N tonne.yr^-1^ and 8.38 P tonne.yr^-1^) and brown noddies (8.43 N tonne.yr^-1^ and 8.38 P tonne.yr^-1^ (dry weight; Table S1). Total N and P in fresh seabird droppings was similar for the three species, with total N ranging from 10.4% (brown noddy) to 11.1% (sooty tern) and total P ranging from 8.2% (sooty tern) to 10.6% (red-footed booby; Table 1). For micronutrients, brown noddy droppings had the highest quantities of Fe and Cu (330 μg.g^-1^ and 29.2 μg.g^-1^ dry weight, respectively), while red-footed booby droppings had the highest Zn and Mg (319 μg.g^-1^ and 16.8 μg.g^-1^ dry weight, respectively; Table 1).

**Table 1.**
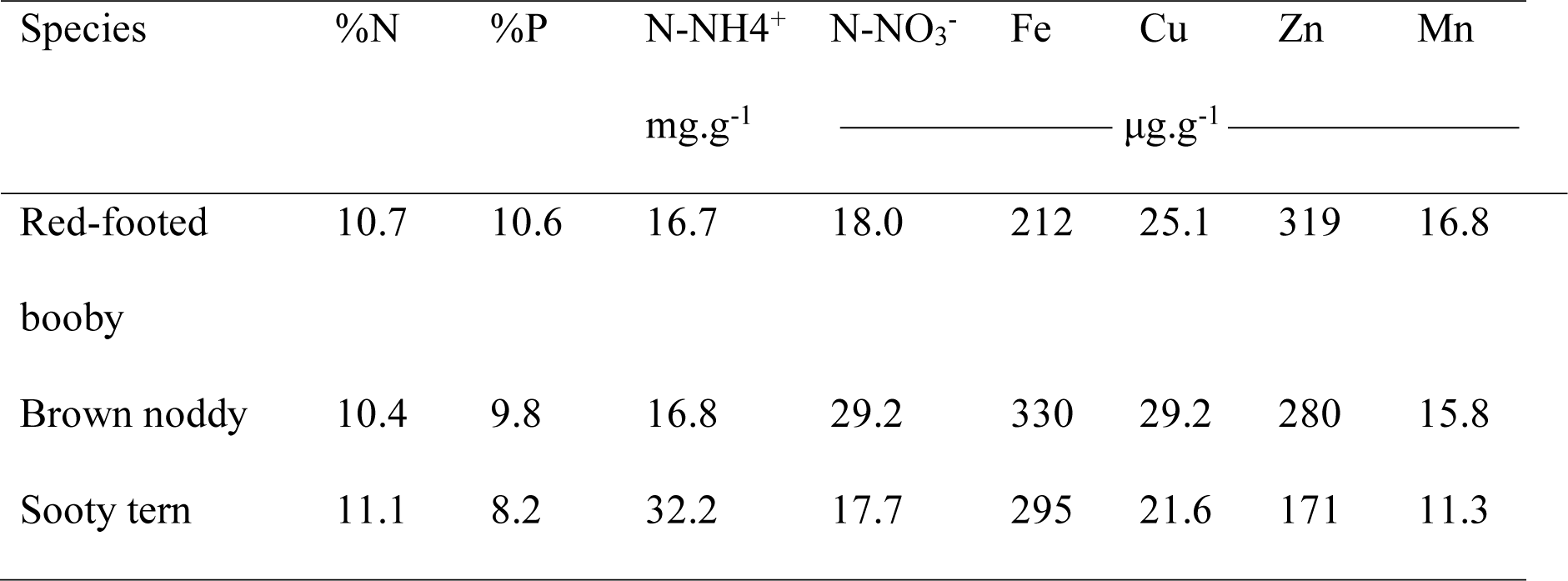
Nutrient concentrations (total nitrogen [%N], total phosphorus [%P], ammonium [N-NH ^+^], nitrate [N-N0 ^-^]) and trace elements (iron [Fe], copper [Cu], zinc [Zn], manganese [Mn]) in seabird droppings. Concentrations are given in dry weight.

### 3.2. Seabird-derived nutrient transfer across taxa and seasons

Sooty tern guano δ^15^N was higher than for brown noddy (adjusted *p* = 0.04; Table 2). Soil and seagrass δ^15^N were higher in the red-footed booby and tern colonies compared to the control for both wet and dry seasons (Figure 2a,b; Table S2). For coastal plants, δ^15^N in the red-footed booby colony was higher than the control during both seasons (wet: adjusted *p* = 0.04, dry: adjusted *p* = 0.03), but for the tern colony δ^15^N was higher than the control during the wet season only (adjusted *p* = 0.02; Figure 2b). Coastal vegetation δ^15^N in the tern colony was lower during the dry than the wet season (adjusted *p* = 0.002).

**Figure 2.**
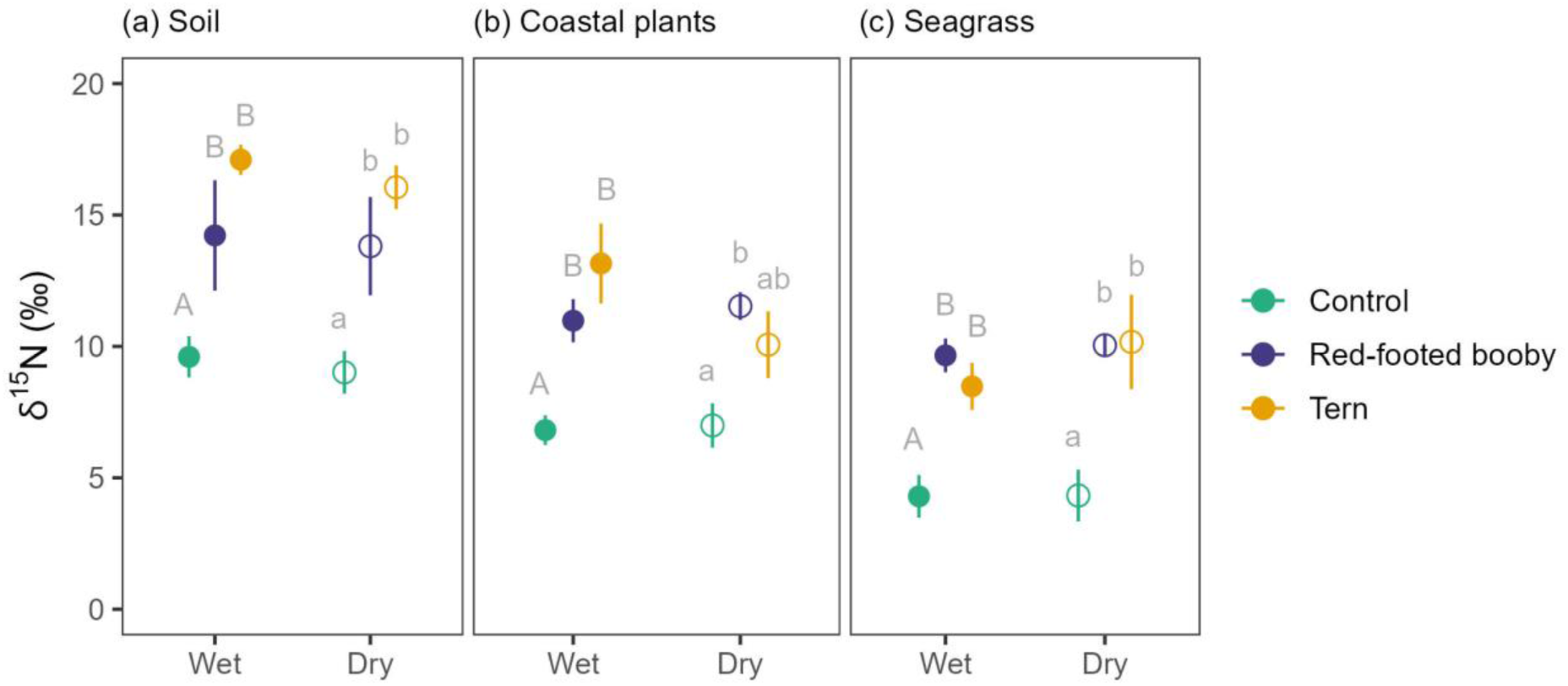
δ^15^N ‰ (Mean ± *SE*) of each treatment (control, red-footed booby, tern) in wet and dry seasons, for soil (a), terrestrial coastal plants (b) and seagrass (b). Closed and open circles represent the wet and dry season, respectively. Uppercase and lowercase letters denote significant (p ≤ 0.05) differences between means for the wet and dry seasons, respectively. Different letters are significantly different.

**Table 2.** Overall range, mean ± *SD* of δ^15^N values for each ecosystem component sampled across the study period and sampling locations.

| Component | Species | n | range (‰) | Mean ± *SD* (‰) |
| --- | --- | --- | --- | --- |
| Guano | *Sula sula* | 28 | 6.41 – 21.9 | 10.8 ± 3.40 |
| | *Anous stolidus* | 25 | 7.64 – 15.0 | 9.77 ± 1.56 |
| | *Onychoprion fuscata* | 25 | 8.76 – 13.2 | 10.6 ± 1.12 |
| Soil | | 36 | 7.08 – 20.4 | 13.3 ± 4.27 |
| Plant | *Tournefortia argentea* | 24 | 4.31 – 13.9 | 9.24 ± 2.91 |
| | *Achyranthes aspera* | 12 | 7.20 – 17.6 | 11.3 ± 3.62 |
| Seagrass | *Thalassodendron ciliatum* | 36 | 0.35 – 17.0 | 7.83 ± 3.47 |

### 3.3. Nutrient levels in terrestrial and nearshore habitats

Soil samples all had a pH of 7.0–9.0 (Table S3), with high variation in EC (0.57 ± 0.77, Mean ± *SD*, n = 36) and CEC (12.1 ± 10.4, Mean ± *SD*, n = 36). Within the cationic exchange complex, Ca^2+^ was the most dominant cation, followed by Mg^2+^, Na^+^ and K^+^ (Table S3). The first two axes of the principal component analysis explained 84% of the variance of the whole dataset (Figure 3), with axis 1 segregating roughly the tern colony from the red-footed booby and control groups, and axis 2 tending to separate the red-footed booby colony from the control group, especially for dry season samples. Soil physical and chemical parameters differed between groups, with higher nutrient content in the tern colony (PERMANOVA tern vs control: adjusted *p* = 0.05, tern vs red-footed booby: adjusted *p* = 0.03). Soil nutrient content was lower in wet season compared to dry season (PERMANOVA *F*_1,35_ = 7.01, *p* = 0.002; Figure 3). Soil pH was negatively correlated to all other parameters indicating soil acidity increased with increasing nutrients.

**Figure 3.**
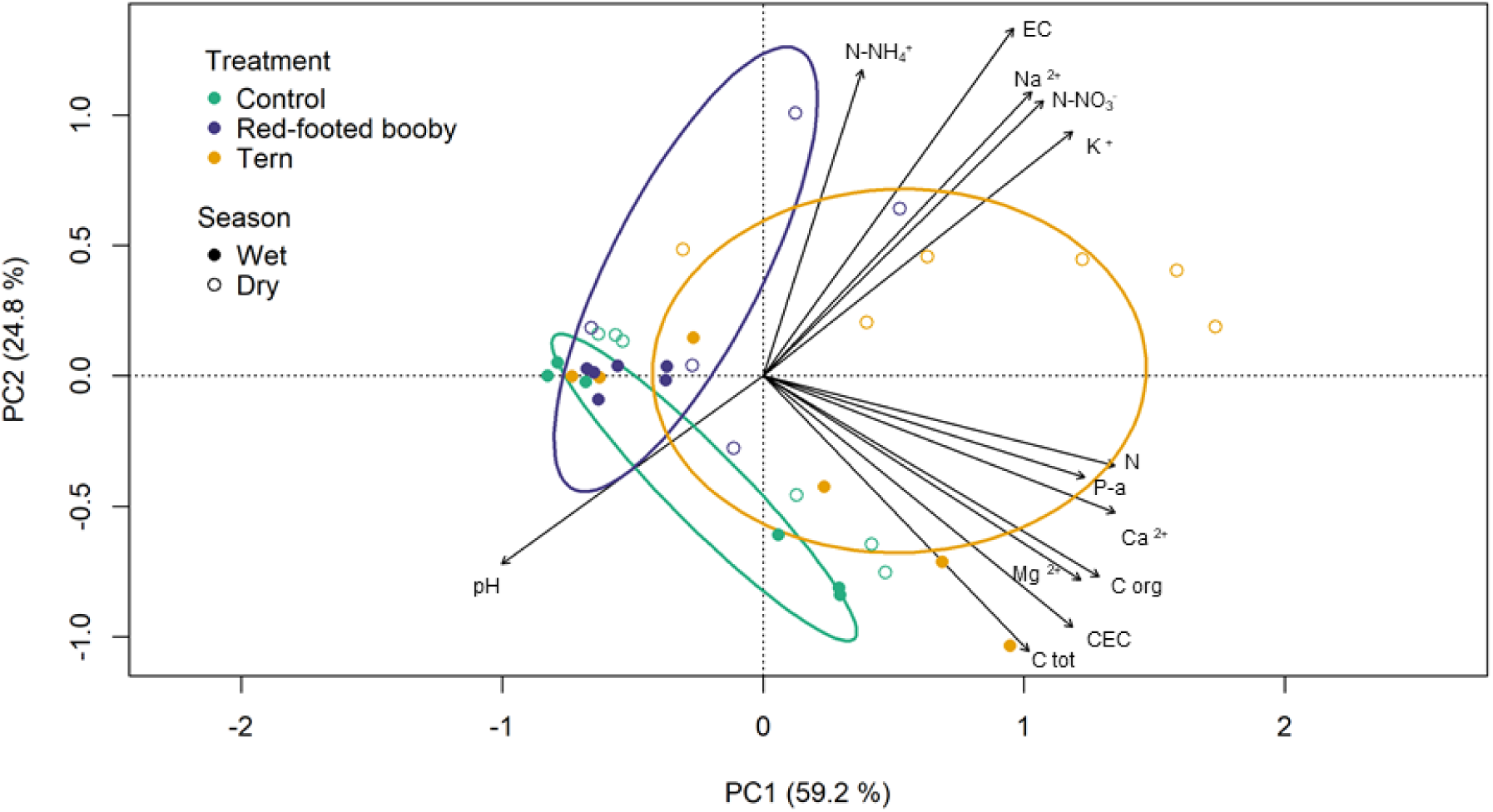
Results from principal component analysis (PCA) on the relationships between soil physical and chemical parameters across treatment and season. Points represent transect observations, with colour depicting the treatment group and shape depicting the season. Black arrows represent the strength (arrow length) and direction of soil variable gradients. Ellipses represent 1 *SD* from the centroid of each treatment group.

For coastal plants, foliar %N was higher in the red-footed booby and tern colonies compared to the control site during the wet and dry seasons (Figure 4a; Table S4). Foliar %N was lower in the dry season than the wet season for the control group (adjusted *p* = 0.01). Leaf %P did not differ between groups in either season, but each group showed a decrease in %P from the wet to the dry season (control: adjusted *p* = 0.004, red-footed booby: adjusted *p* = 0.01, tern: adjusted *p* = 0.003; Figure 4b).

**Figure 4.**
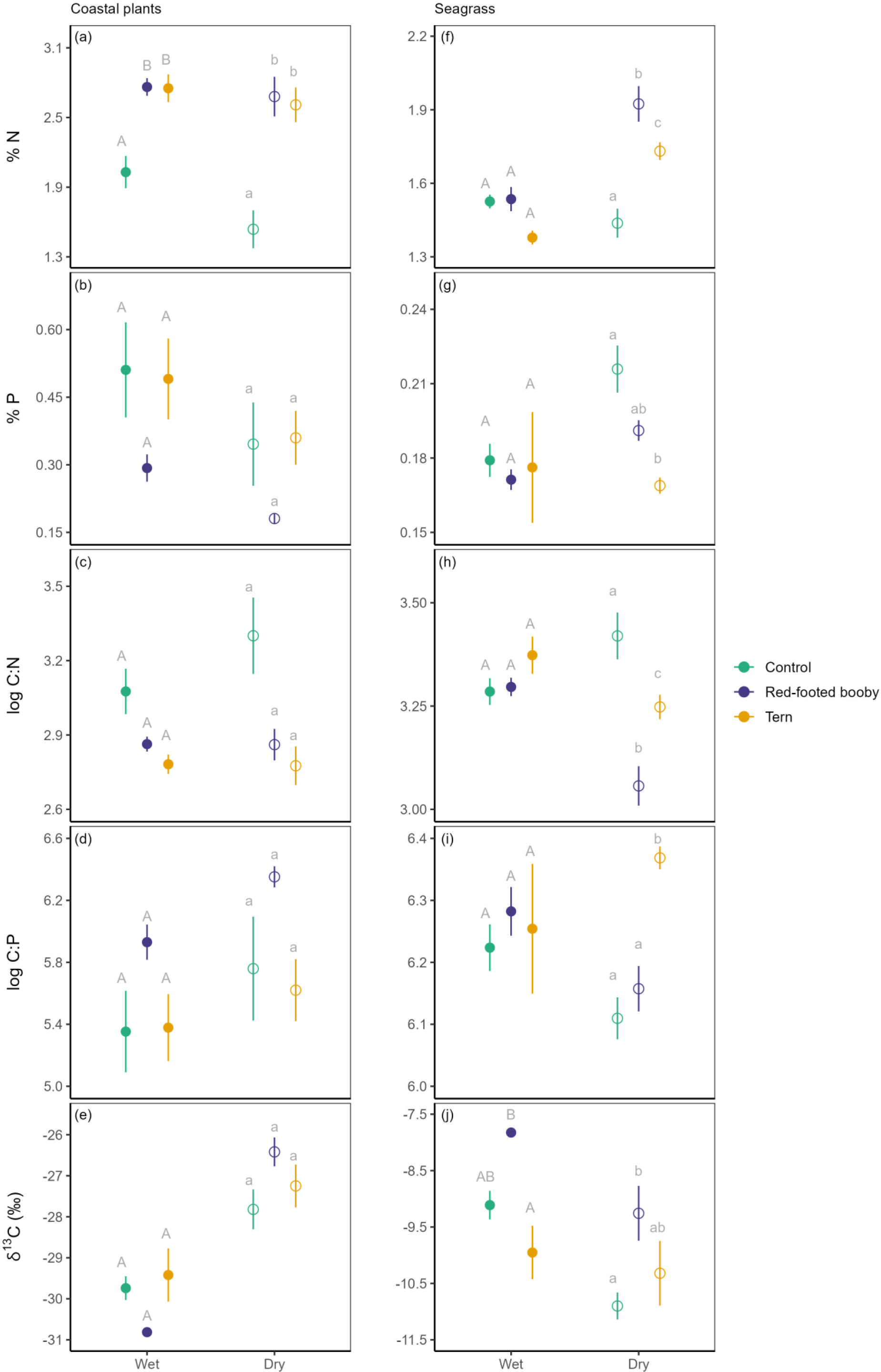
Mean (± *SE*) of nutrient levels (expressed in % dry weight), log nutrient ratios and δ^13^C ‰ of each group (control, red-footed booby, tern) in wet and dry seasons, for terrestrial coastal plants (a-e) and seagrass (f-j). Closed and open circles represent the wet and dry season, respectively. Uppercase and lowercase letters denote significant (p ≤ 0.05) differences between means for the wet and dry seasons, respectively. Different letters are significantly different. Note differences in y-axis scales. Error bars absent in some cases due to scaling.

For seagrass, foliar %N was higher in the red-footed booby and tern colonies compared to the control during the dry season only (Figure 4f). Seagrass %N was higher in the dry season compared to the wet season for the red-footed booby (adjusted *p* < 0.0001) and tern group (adjusted *p* < 0.0001). %P in seagrass leaves was lower closer to the tern colony compared to the control site in the dry season (adjusted *p* = 0.01; Figure 4g). Seagrass %P was higher during the dry season than the wet season for the control group (adjusted *p* = 0.03; Table S5). There was a linear increase in foliar %N with increasing foliar δ^15^N for coastal plants during the wet and dry seasons, and for seagrass during the dry season only (Figure 5).

**Figure 5.**
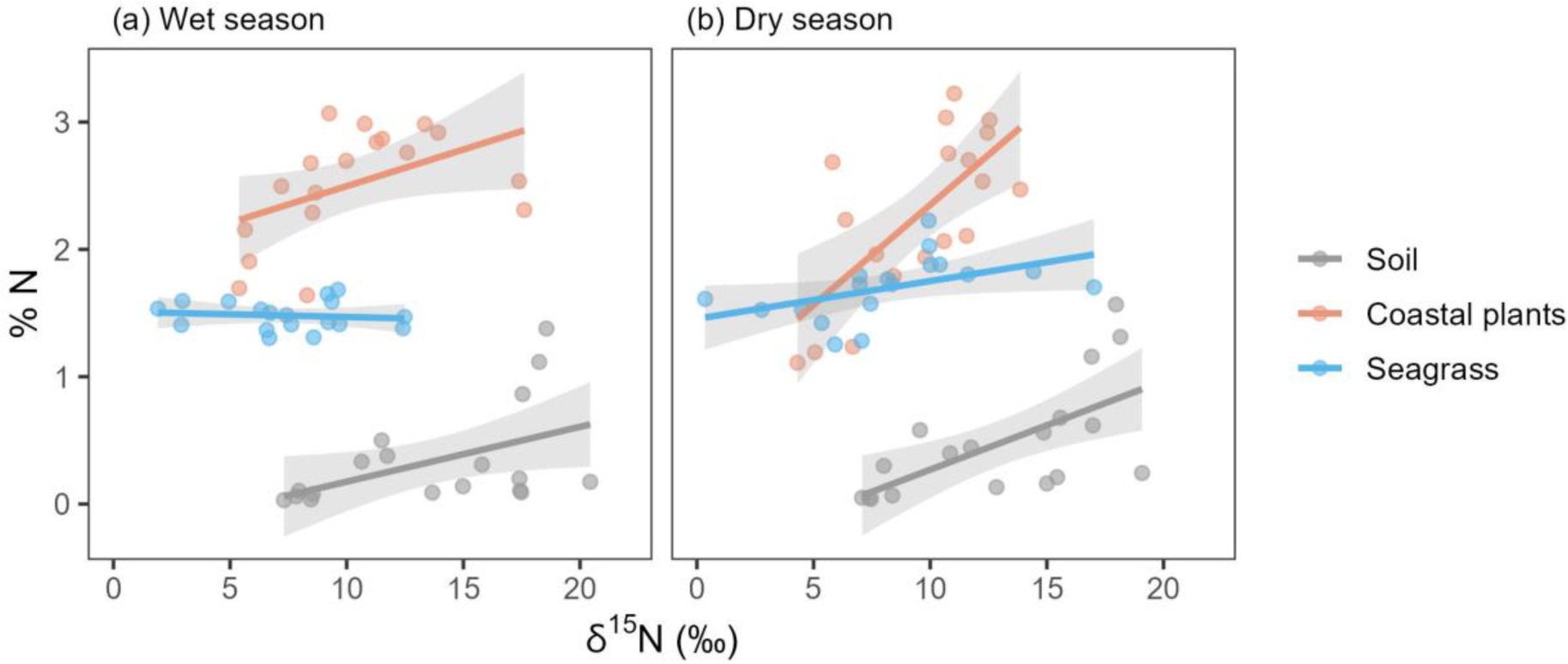
Relationships between δ^15^N and %N during the wet (a) and dry (b) season for soil, coastal plants and seagrass. Shaded areas represent 95 % CI.

### 3.4. Coastal and marine plant nutrient limitations and growth rates

For coastal plants, log C:P ratios increased from wet to dry seasons for all groups (control: adjusted *p* = 0.0008, red-footed booby: adjusted *p* = 0.0005, tern: adjusted *p* = 0.03; Figure 4d, Table S4). For seagrass, log C:N values differed between all groups during the dry season (Figure 4h; Table S5) and between the wet and dry season for all groups (control: adjusted *p* = 0.01, red-footed booby: adjusted *p* = 0.0001, tern: adjusted *p* = 0.02). Log C:P ratios were higher in the tern colony during the dry season (control vs tern: adjusted *p* = 0.005, tern vs red-footed booby: adjusted *p* = 0.02; Figure 4i).

Leaf δ^13^C in coastal vegetation did not differ between groups in either season, but rather increased from the wet to the dry season for all groups (Figure 4e; Table S4). For seagrass, δ^13^C was higher near the red-footed booby colony compared to the tern colony during the wet season (adjusted *p* = 0.002) and higher near the red-footed booby colony than the control during the dry season (adjusted *p* = 0.02; Figure 4j). Furthermore, δ^13^C in seagrass decreased from the wet to the dry season near the control (adjusted *p* = 0.0006) and red-footed booby colony (adjusted *p* = 0.003).

## DISCUSSION

Our study shows that seabird nutrient contributions to atolls can be substantial, with a combined total from all three seabird species of 64.5 N t.yr^-1^ and 49.1 P t.yr^-1^ estimated in our study system. Breeding sooty terns contributed the highest quantities of nutrients, coinciding with the dry season. We detected persistent seabird-derived nutrient transfer from red-footed booby and tern colonies to soil, coastal plants and seagrass. Soil macro- and inorganic nutrient levels were higher in the tern colony during the dry season, while foliar nitrogen in coastal plants was higher in both seabird colonies year-round. Seabird-derived nutrients reversed nitrogen limitation of seagrass during the dry season.

### Seabird nutrient contributions across taxa

Population size is a key driver in nutrient quantity since it dictates the number of individuals acting as nutrient vectors (Subalusky & Post, 2019). In our study, sooty terns comprised the largest population and contributed the highest quantities of nutrients. Our estimated nutrient inputs align with those of agriculture practices (60 – 400 N kg.ha^-1^.yr^-1^ Young et al., 2010; 156 N kg.ha^-1^.yr^-1^ and 30 kg P kg.ha^-1^.yr^-1^ McFadden et al., 2016), and other tropical seabird islands (this study: 151.4 N kg.ha^-1^.yr^-1^, 115.3 P kg.ha^-1^.yr^-1^; Heron Island: 324.1 N kg.ha^-1^.yr^-1^, 65.5 P kg.ha^-1^.yr^-1^ [Staunton Smith & Johnson, 1995]; Palmyra Atoll: 687 N kg. ha^-1^.yr^-1^ and 111 P kg.ha^-1^.yr^-1^ [Young et al., 2010]). However, our estimates do not account for nutrient contributions from non-breeding seabirds, which can constitute a large proportion of total populations (Schreiber & Chovan, 1986); for example, non-breeders can comprise 33% of breeding populations and spend 50% less time than breeders in their colony in temperate areas (Riddick et al., 2012). Quantification of non-breeding individuals year-round and their diurnal movement patterns would refine our estimates.

Tropical seabirds exhibit overlapping diets with a large diversity of prey items (Cherel et al., 2008; Catry et al., 2009). Consequently, guano macronutrient levels were similar for our three study species. Micronutrient content in guano concurs with that of their prey, comprising small pelagic fish and cephalopods rich in iron and zinc (Sabino et al., 2022). These concentrations are consistent with results of recent reviews and meta-analyses, e.g. Suliforme droppings contain the highest levels of phosphorus and zinc compared to other seabird orders (De La Peña-Lastra, 2021a; Grant et al., 2022). Guano δ^15^N for sooty terns was slightly higher than for brown noddies, which may be due to differences in metabolism and the main fish species consumed (De La Peña-Lastra, 2021a). In the Western Indian Ocean, sooty terns and brown noddies diet includes post-larval goatfish, anchovies and flying fish (Jaquemet et al., 2008; Catry et al., 2009). Here, sooty terns and brown noddies nested on the same island and their guano δ^15^N did not differ from that of red-footed boobies, indicating similar nutrient isotopic baselines for these two seabird islands.

### Seabird-derived nutrient transfer

In the red-footed booby colony, the high δ^15^N in both seasons indicated persistent transfer of seabird-derived nutrients to terrestrial and nearshore marine habitats linked to their year-round breeding patterns. In contrast, despite the tern colony breeding exclusively in the dry season, we also detected high δ^15^N during the wet season in terrestrial and nearshore ecosystems. This is presumably linked partly to brown noddies breeding in much smaller numbers during the wet season (ICS, unpubl. data) and individuals which can remain at their breeding grounds year-round (Bailey, 1968; Lebarbenchon et al., 2023). However, because of isotopic fractionation of guano, nutrient input during the breeding season can maintain isotope levels throughout the non-breeding season (Pascoe et al., 2022). High δ^15^N values in sediment have also played an important role in identifying abandoned seabird colonies (Mizutani et al., 1991) and tracking seabird activity over centuries from sediment cores (Michelutti et al., 2009).

### Nutrient levels in terrestrial and nearshore habitats

Although both seabird colonies had higher soil nutrient concentrations than the control site, soil macro- and inorganic nutrient levels were higher in the tern colony than in the red-footed booby colony. This is linked to the higher nest density of terns (Table S1) and nesting location (Smith et al., 2011). Nutrients from tree-nesters, like red-footed boobies, are more sparsely distributed and less likely to reach the soil compared to ground-nesters (Gaiotto et al., 2022). Furthermore, the red-footed booby colony’s presence on South Island is relatively recent (ICS, unpubl. data), in contrast to the long-established tern colony, where nutrients have accumulated over several decades. However, observed patterns may also be due to island size since the tern colony is on the smallest island of the study. On small islands, subsidy impacts are more pronounced since they have a greater perimeter-to-area ratio which enable greater per-unit-area effects (Pascoe et al., 2021; Obrist et al., 2022). Furthermore, soil nutrient levels were lower during the wet season, likely due to rainfall which increases solubility and loss of soil nutrients, either by plant uptake, or through leaching and surface run-off (De La Peña-Lastra, 2021a). However, in the tern colony, higher nutrients during the dry season are almost certainly related to breeding seasonality. The relatively short occurrence of very high numbers of breeding sooty terns and brown noddies during the dry season leads to an annual pulse in nutrient resources. Year-round sampling would elucidate the duration of effects and nutrient dynamics of this resource pulse. Additional studies can also assess intra-island nutrient variations, by documenting nest densities at sampling sites for within-colony comparisons, or at multiple distances outside the colony (Pascoe et al., 2022).

The persistent δ^15^N enrichment in the seabird colonies led to year-round biologically relevant uptake of nitrogen (%N) in coastal vegetation. Average foliar %N was 1.4–1.7 times higher in both seabird colonies than the control site in both seasons. In contrast, foliar %P showed no response to seabird contributions, which has been observed elsewhere (Mulder et al., 2011). Phosphorus is less soluble than nitrogen and strongly adsorbed in soils, especially the calcareous soils of atolls which precipitate phosphorus as calcium phosphate (Otero et al., 2015). Phosphorus stability in soils can be long-lasting, even reflecting seabird-derived phosphorus from extinct colonies (Wardle et al., 2009; De La Peña-Lastra et al., 2021b; Mutillod et al., 2023). Red-footed boobies historically bred in large numbers on our control island, but not since at least 1897 (Feare, 1978), suggesting coastal plants on this island may derive phosphorus from long-term reserves (Mulder et al., 2011). Moreover, lower rainfall facilitates phosphorus retention in soil, explaining the marked decrease in foliar %P during the dry season in all groups.

Despite the year-round transfer of seabird-derived nutrients to seagrass in the nearshore environment, δ^15^N enrichment resulted in biologically relevant nitrogen uptake (seagrass foliar %N) at the seabird colonies only during the dry season. This was contrary to our expectations, since higher rainfall increases nutrient loadings in nearshore environments through surface run-off or leaching (Rankin & Jones, 2021). The higher biomass of seabirds during the dry season suggests that a critical threshold in seabird density is needed to generate responses in seagrass foliar nitrogen. Similar results were shown in Baltic Sea islands, with high nitrogen values in marine algae were detected only at islands with high seabird nesting densities (Kolb et al., 2010).

The carbonate substrate of tropical seagrass binds strongly with phosphorus, maintaining phosphorus levels in seagrass, even after many decades of discontinued seabird contributions (Herbert & Fourqurean, 2008), allowing us to make similar conclusions as with coastal plants. The dry season decrease in seagrass phosphorus levels near the tern colony is likely related to seagrass location and local oceanographic conditions. Seagrass near the tern colony is less sheltered than seagrass growing in the lagoon near the other two treatment groups. Exposed areas have lower rates of sediment oxygen uptake and consequently, lower rates of sediment phosphorus release compared to sheltered sites (Jensen et al., 1998; Fourqurean & Zieman, 2002). During the dry season, increased wave activity by the south-east monsoon winds may further reduce this process in seagrass at the tern colony, leading to decreased phosphorus availability.

### Coastal and marine plant nutrient limitation and growth rates

Although seabird-derived nutrients increased foliar %N in coastal plants, we did not detect any reductions in nutrient limitation (log C:N, log C:P) to growth. This implies that other factors limit coastal plants in their responses to seabird contributions on atolls, such as limited availability of other essential macro- and micro-nutrients (Lavelle et al., 2005). Additionally, we did not find differences in foliar δ^13^C. In temperate locations, positive correlations between higher foliar seabird-derived nutrient levels and δ^13^C enrichment have been documented, suggesting faster plant growth (Wainright et al., 1998). The lack of response in our study could be due to low sample sizes. We did detect increases in foliar δ^13^C from the wet to the dry season, confirming water stress in the latter. To reduce water loss, plants reduce stomatal conductance, leading to reduced discrimination of ^13^C isotopes and causing δ^13^C to increase (Mulder et al., 2011).

In nearshore environments, response to seabird-derived nutrients is also governed by local geomorphological and oceanographic features, such as bottom topography, wave action, currents and depth (Kazama, 2020; Signa et al., 2021). For seagrass, other factors such as epiphyte load, light availability and grazing pressure all interact to influence their functioning (Frankovich & Fourqurean, 1997; Anderson & Fourqurean, 2003). In our study, the transfer and uptake of seabird-derived nitrogen in seagrass during the dry season led to reduced nitrogen limitation (log C:N), coinciding with the higher biomass of breeding seabirds during this season. However, a larger sample size and quantification of seagrass physical parameters is needed to clarify the response of seagrass to seabird contributions at our study site.

### Conservation implications

From our results we can draw recommendations for management of seabird populations on atolls. Nutrient input was driven by sooty terns in our study because of their higher biomass and ground-nesting habits. Sooty terns are the most abundant tropical seabird (Hughes et al., 2017), therefore maintaining or restoring their populations and other high-density ground-nesting seabirds should be prioritised (e.g. with invasive alien mammal eradications, assisted colonisation, management of breeding habitat), since they have the greatest potential for supporting and boosting resilience of atolls and other tropical island ecosystems (Berr et al., 2023). In Seychelles, sooty tern eggs have been exploited commercially since the beginning of the 20^th^ century (Hutchinson, 1950; Feare, 1976a). Managed by local authorities, harvesting strategies are largely based on population demographic parameters (Feare, 1976b; Feare & Doherty, 2004, 2011). By quantifying sooty tern nutrient contributions of an unharvested colony on Farquhar Atoll, we highlight an additional critical element to consider in harvesting decisions. Reducing harvesting quotas and island colonies exploited, or replacing harvesting with sustainable ecotourism practices such as birdwatching, will reduce disturbance to sooty tern colonies.

Our results also show that other tropical seabirds, e.g. red-footed boobies, supply nutrients continuously to atolls because of their lengthy breeding and roosting patterns, which helps maintain nutrient status of soil, plants and nearshore marine communities. Seabird conservation efforts should therefore also target species that breed year-round. Ultimately restoring, and supporting breeding populations of a range of seabirds on islands is likely to provide the widest range of nutrients throughout the year and contribute the most to atoll ecosystem functioning (Benkwitt et al., 2022).

## Conclusion

We present the first research on seabird contributions to atoll ecosystems in Seychelles, showing the spatial and temporal influences that tropical seabird subsidies have on soil and plants, and seagrass growing adjacent to atolls. Our study builds on research showing that response to seabird contributions is influenced by seabird biomass, breeding and foraging behaviours, seasonality and solubility of nutrients (Smith et al., 2011). We quantify and highlight the significance of sooty tern nutrient contributions to the maintenance of atoll ecosystems. Seabird-subsidised atolls play substantial roles in maintaining biodiversity and marine connectivity. Our research provides insights into in a relatively undisturbed system which can be used as a benchmark site with which to compare more directly impacted atolls and future changes.

## ACKNOWLEDGEMENTS

We thank Island Conservation Society for providing logistical support during the fieldwork, and Islands Development Company for travel and permission to work on Farquhar Atoll. Funding was provided by the Rufford Foundation, the University of Reunion Island International Relations Department and the Bertarelli Foundation through the Marine Science Programme. JA was supported by a doctoral fellowship from the Reunion Island Regional Council. Fieldwork was conducted under permit number A0157 by Seychelles Bureau of Standards.

## AUTHOR CONTRIBUTIONS

Conceptualization: J.A, N.B, S.J. Investigation and data curation: J.A, J.L, A.H. A.G. Methodology, formal analysis, writing - original draft: J.A. Writing - review and editing: N.B, J.L, A.H, N.A.J.G, G.R, M.L.C, S.J.

## DATA AVAILABILITY STATEMENT

The data that support the findings of this study will be available in Dryad Digital Repository upon acceptance.

