## Supplementary methods, Table S1-S5 for "Seabird traits and seasonality modulate nutrient dynamics of terrestrial and marine habitats on atolls"

**1. Study area**

Farquhar Atoll (10°11′ S, 51°06′ E) is the most southerly atoll of the Seychelles archipelago in the Western Indian Ocean. Farquhar lies ca. 770 km from Mahé, the main island of Seychelles, and 285 km northeast of Madagascar (Figure 1). It is a low-lying, triangular-shaped atoll (total area 178 km^2^), consisting of 11 islands (total landmass 8 km^2^) surrounding a shallow lagoon (Duvat et al., 2017). The majority of Farquhar’s landmass consists of North Island (3.32 km^2^), South Island (3.69 km^2^) and Goëlettes (0.32 km^2^). Anthropogenic activity on Farquhar is limited to a small settlement on North Island managed by Islands Development Company. The settlement supports an ecotourism establishment based on a seasonal catch and release fly fishing activity operated by Blue Safaris Seychelles. The environmental NGO Island Conservation Society is also present on the island.

On North and South Islands, the vegetation cover along the beach margin of the ocean and lagoon shores consists of native coastal shrubs including *Scaevola taccada*, *Tournefortia argentea* and *Suriana maritima*. Moving inland, the vegetation transitions into a mix of indigenous and introduced grasses, sedges and trees. The interior of the islands has been heavily altered by historical exploitation of copra and timber, and is covered in introduced *Cocos nucifera* and *Casuarina equisetifolia* (Stoddart & Poore, 1970). Distinctively, three tidal swamps enclosed by sand bars can be found on South Island’s lagoon shore, which are dominated by *Pemphis acidula*. In contrast, Goëlettes is treeless and almost entirely covered in a low herb plant community comprised of *Boerhavia repens*, *Achyranthes aspera*, grasses and sedges, and with some coastal shrubs along its lagoon shore (Stoddart & Poore, 1970). Adjacent to island shores in the lagoon and on the reef flats, there are large expanses of seagrass, dominated by *Thalassodendron ciliatum* (Stokes et al., 2019).

Farquhar is home to large breeding colonies of red-footed boobies *Sula sula*, brown noddies *Anous stolidous* and sooty terns *Onychoprion fuscatus*, on separate islands (Duhec et al., 2017). Sooty terns and brown noddies nests on Goëlettes, estimated at 208,625 and 19,139 breeding pairs respectively (Table S1; ICS unpubl. data). Sooty terns and brown noddies are similar in size (140‒240 g and 160‒205 g, respectively; Schreiber & Burger, 2001) and form simple nests, consisting of a shallow depression on the ground. Brown noddies breed mainly between May and October and in much lower numbers throughout the year, whereas sooty terns only breed between May and October. Two large red-footed booby colonies are located in the tidal swamps and along the lagoon coastline of South Island, estimated at a total of 10,228 breeding pairs annually (Figure 1, Table S1; ICS unpubl. data). Red-footed boobies are heavier (800‒1500 g; Schreiber & Burger, 2001) and breed year-round with peaks in March-April and November-December. They build nests around 1‒2 m from the ground in *P. acidula* and along the lagoon shore in *T. argentea*. All three species are surface pelagic feeders, preying mainly on fish and cephalopods (Weimerskirch et al., 2005; Catry et al., 2009). Other seabirds breed on Farquhar’s islets but in relatively much small numbers (< 100 breeding pairs each), including black-naped terns *Sterna sumatrana*, roseate terns *Sterna dougallii*, lesser noddies *Anous tenuirostris*, greater-crested tern *Thalasseus bergii* and fairy tern *Gygis alba* (Duhec et al., 2017). Due to these attributes, Farquhar's islets are designated as Important Bird Area by BirdLife International (Rocamora & Skerrett, 2001). North Island, in contrast, has very few breeding seabirds, attributed to the islands’ historical use the main centre for human settlement and coconut exploitation (Duhec et al., 2017). Most rain on Farquhar fall between November and April as a result of north-east monsoon winds. Between May and October, trade winds blowing from the south-east result in lower rainfall (Piggot, 1961).

**2. Sampling design**

We investigated the influence of seabird traits on nutrient dynamics using three separate islands and their breeding seabird species as a treatment group: (a) red-footed boobies on South Island, (b) terns, comprising brown noddies and sooty terns, on Goëlettes, and (c) North Island, with no breeding seabirds, as a control island. Because of their relatively low numbers, we did not account for additional breeding species on Goëlettes (see above) and assumed they make a relatively small contribution to seabird-derived nutrient dynamics. Because rainfall influences nutrient dissipation and nutrient availability in nearshore environment (Signa et al., 2021), we incorporated local seasonality by sampling in both the wet (March 2022) and dry season (August 2022).

**3. Sampling of seabird droppings**

We collected fresh seabird droppings of the three breeding species within their colonies to determine nutrient concentrations. For red-footed boobies, guano was collected on a black plastic sheet (2 x 2 m) pinned to the ground below nesting and roosting individuals. Guano was sampled shortly after deposition on the first day, with additional droppings obtained by leaving the plastic sheet overnight and collected in the early morning. For brown noddies and sooty terns, we collected droppings from fresh samples on vegetation near breeding and roosting individuals. Individual droppings were combined to obtain a minimum of 30 g wet weight per species for nutrient analyses (Staunton Smith & Johnson, 1995). Separate samples were collected for isotopic analysis. All samples were kept cool in the field and stored frozen until further processing.

We determined macro- and micro-nutrient content of fresh seabird droppings for each species. Nitrogen (N) was determined according to the Kjeldahl method (Kjeldahl, 1883) (Büchi KjelMaster K-375, Switzerland). Ammonium (N-NH_4_^+^) was assayed by distillation and titration with sulphuric acid, while nitrate (N-N0_3_^-^) was extracted in water with volume ratio of 1:4 and assayed by continuous flow colorimetry (Proxima, Alliance Instrument, USA). Phosphorus (P) and micro-nutrients were determined by preliminary dry combustion (600 °C), then dissolving the residual material in 6 M HCl. P concentration was obtained by extraction using ammonium molybdate and ascorbic acid and measured with a continuous flow colorimeter, while concentration of iron (Fe), manganese (Mn), zinc (Zn) and copper (Cu) were determined by atomic absorption spectrophotometry (AAnalyst 400 spectrometer, PerkinElmer, USA).

We estimated the total annual N and P input from seabird droppings for each species based on previously used methods (Riddick et al., 2018; Graham et al., 2018).

$${Nutrient}_{gi}=Q_{gi}\times{Dr}_{i}\times\left[ \left( {Bd}_{i}\times T_{i}\times f_{i} \right)+\left( \frac{P_{i}}{2}\times{Bd}_{i} \right) \right]$$

Annual input per nutrient type and species (${Nutrient}_{gi}$, t.yr^-1^) was obtained from the quantity of N or P measured in seabird droppings for each species ($Q_{gi}$, mg.g^-1^), the excrement rate of the species (${Dr}_{i}$, g.bird^-1^.day^-1^), the number of breeding adults for that species (${Bd}_{i}$), the length of the breeding period (number of days from courtship to end of chick-rearing, $T_{i}$), the proportion of time spent at the colony during breeding (to account for absence of birds during feeding bouts, $f_{i}$) and the productivity of the species (fledged chicks per breeding pair, $P_{i}$). The excrement rate for red-footed boobies was obtained from Young et al., 2010. For brown noddies and sooty terns, we estimated excrement rates based on measurements for black noddy *A. minutus* scaled allometrically to body size according to Staunton Smith & Johnson, 1995. Values for the length of the breeding period, proportion of time at the colony and productivity for each species were obtained from a literature review by Riddick et al., 2012. On Farquhar, red-footed booby and brown noddy colonies also contain roosting individuals, while sooty tern colonies do not. We lacked data on non-breeders so our estimates are for breeding seabirds only.

**4. Terrestrial and marine sampling**

We sampled soil and terrestrial coastal vegetation in two different habitat types on each island; coastal shrub habitat and low herbaceous habitat. In each habitat type, samples were collected along three randomly placed 50-m transects set a minimum 100 m apart and parallel to the lagoon shore. As South Island had no low herbaceous habitat, coastal shrub habitat was sampled at each of the two red-footed booby colonies (Figure 1). On the two seabird islands, we sampled within the colonies. Along each transect we collected soil within the first 5 cm at three evenly spaced positions, then homogenized to obtain a composite surface soil sample. Soil was sieved (< 2 mm mesh) to remove rocks and debris (Young et al., 2010). A sub-sample was obtained for isotope analysis, while the remaining was used for nutrient analyses. Along the transect, we collected samples of green leaves in full sun from five individuals of *T. argentea* in the coastal shrub habitat and *A. aspera* in the low herb habitat. For each transect, we sub-sampled one leaf from each plant individual for isotope analysis and the remaining leaves were homogenized for nutrient analyses.

For the marine sampling, we sampled the seagrass *T. ciliatum* adjacent to each island. Sampling locations were at least 100 m apart and < 300 m from the island. We collected seagrass leaves at six random locations within a radius of 5 m. A sub-sample from each location was obtained for isotope analysis. All samples for isotope analysis were stored frozen upon return from the field until further processing, and the remaining samples were air-dried. We sampled the same locations in both seasons.

**5. Nutrient analyses**

Upon collection, plant material was rinsed with freshwater, dried to a constant mass in a drying-oven and powdered using a ball mill. Soil was equally dried in an oven and soil pH were measured in a soil:water ratio of 1:5 (PHM220 pH meter, Radiometer analytical, France) and electrical conductivity (EC) was measured in a 1:5 soil:water suspension (CDM 210 conductivity meter, Radiometer analytical, France). Total carbon (C) for soil and terrestrial plant samples was determined using an element analyser (Vario Max Cube, Germany). Organic carbon (C org) for soil and seagrass, and total N for soil and plant samples were determined using the Dumas method (Black et al., 1965) with an element analyser. Soil N-NH_4_^+^ and N-NO_3_^-^ were determined in a 1:5 soil extract with 1 M KCl solution by continuous flux colorimetry. Bioavailable phosphorus (P-a) in soil was determined by the Olsen-Dabin method (Dabin, 1967) using 0.5 M sodium hydrogen carbonate solution and a continuous flow colorimeter. Soil cationic exchange capacity (CEC) and exchangeable cations (Ca^2+^, Mg^2+^, Na^+^ and K^+^) were determined by hexamine cobalt chloride extraction (Ciesielski et al., 1997) and assayed by atomic absorption spectrophotometry. Total P in plant samples was analysed similarly to seabird droppings. Sample blanks and reference materials were analysed within each analytical run. Nutrient analyses were conducted at the CIRAD-Reunion in the laboratory of agronomic analyses.

**6. Isotopic analyses**

Sub-samples of seabird guano, soil and plant material were dried in a drying oven at 50°C for minimum of 48 h and powdered using a ball mill. All samples were combusted using an Elementar Vario Micro Cube Elemental Analyser and δ^15^N and δ^13^C were measured using an Isoprime 100 Isotope Ratio Mass Spectrometer with international standards IAEA 600, USGS 41 and CH6, at the stable isotope facility at Lancaster University (Lancaster, UK). Soil and seagrass samples were run twice, once after repeated acidifications with 1 M HCl to remove carbonates for δ^13^C analyses and once without this treatment for δ^15^N. Accuracy based on internal standards was within 0.2 ‰ standard deviation for δ^15^N and 0.1 ‰ standard deviation for δ^13^C. Selected samples were run in triplicate to further ensure accuracy of readings.

**Table S1.** Island size and seabird characteristics (ICS, *unpublished data*) of the three study islands of Farquhar Atoll, with the amount of total nitrogen and total phosphorus deposited annually by large breeding seabird colonies. na = not applicable.

| Island | Total area (ha) | Main breeding species (number of breeding pairs; year of census) | Colony area (ha) | Nest density (nests.m^-2^) | Seabird biomass (kg.ha^-1^) | N input (t.yr^-1^) | P input (t.yr^-1^) |
| --- | --- | --- | --- | --- | --- | --- | --- |
| North Island | 379 | na |  | na | na | na | na |
| South Island | 394 | Red-footed booby *Sula sula* (10 228; 2021) | 15.3 | 0.07 | 20.7 | 8.43 | 8.38 |
| Goëlettes | 32 | Brown noddy *Anous stolidous* (19 139; 2017) | 2.0 | 0.95 | 48.0 | 6.30 | 5.90 |
|  |  | Sooty tern *Onychoprion fuscata* (208 625; 2021) | 6.72 | 3.10 | 496.8 | 71.19 | 52.19 |

**Table S2.** Results of linear mixed models fitted to test changes in δ^15^N for soil, coastal plants and seagrass under different seabird influence (control, red footed booby, tern) and seasons (wet, dry). Statistically significant results (*p* ≤ 0.05) are indicated in bold. df: Degree of freedom.

| Variable | Random effect | Fixed effects | Mean square | df | F value | P value | Conditional R2 | Marginal R2 |
| --- | --- | --- | --- | --- | --- | --- | --- | --- |
| **Soil** |  | | | | | | | |
| δ^15^N | Sample location | Treatment | 178.570 | 2 | 20.570 | **0.046** | 0.56 | 0.56 |
|  |  | Season | 4.189 | 1 | 0.483 | 0.493 |  |  |
|  |  | Habitat | 54.692 | 1 | 6.300 | 0.129 |  |  |
|  |  | Treatment x Season | 0.329 | 2 | 0.038 | 0.963 |  |  |
| **Terrestrial coastal plant** | | | | | | | | |
| δ^15^N | Sample location | Treatment | 45.666 | 2 | 18.764 | **0.050** | 0.79 | 0.73 |
|  |  | Season | 5.554 | 1 | 2.282 | 0.142 |  |  |
|  |  | Habitat | 39.412 | 1 | 16.194 | 0.057 |  |  |
|  |  | Treatment x Season | 24.106 | 2 | 4.953 | **0.015** |  |  |
| **Seagrass** | | | | | | | | |
| δ^15^N | Sample location | Treatment | 69.356 | 2 | 14.489 | **0.0003** | 0.63 | 0.52 |
|  |  | Season | 4.438 | 1 | 0.927 | 0.351 |  |  |
|  |  | Treatment x Season | 2.291 | 2 | 0.479 | 0.629 |  |  |

**Table S3.** Soil physical and chemical characteristics for each treatment group (control, red-footed booby and tern; n = 12 each group) over the study period. EC = Electrical conductivity, P-a = bioavailable phosphorus, CEC = Cationic Exchange Capacity.

|  | Control | | Red-footed booby | | Tern | |
| --- | --- | --- | --- | --- | --- | --- |
|  | range | Mean ± SD | range | Mean ± SD | range | Mean ± SD |
| pH | 8.05 – 9.49 | 8.48 ± 0.42 | 7.30 – 9.09 | 8.06 ± 0.49 | 7.20 – 9.18 | 7.83 ± 0.61 |
| EC (mS.cm^-1^) | 0.05 – 0.28 | 0.16 ± 0.07 | 0.05 – 2.90 | 0.67 ± 0.97 | 0.07 – 2.10 | 0.88 ± 0.79 |
| Total C (mg.g^-1^) | 115 – 145 | 126 ± 10.3 | 111 – 138 | 121 ± 6.86 | 111 – 143 | 128 ± 9.57 |
| C organic (mg.g^-1^) | 4.71 – 66.2 | 31.7 ± 26.2 | 8.03 – 24.5 | 14.4 ± 5.12 | 7.78 – 100 | 51.6 ± 33.0 |
| Total N (mg.g^-1^) | 0.30 – 5.81 | 2.45 ± 2.12 | 0.61 – 6.18 | 1.92 ± 1.54 | 0.92 – 15.7 | 7.76 ± 5.31 |
| N-NO_3_^-^ (mg.g^-1^) | 0.002 – 0.04 | 0.02 ± 0.02 | 0.006 – 1.15 | 0.27 ± 0.42 | 0.01 – 1.34 | 0.44 ± 0.45 |
| N-NH_4_^+^ (mg.g^-1^) | 0.0008 – 0.003 | 0.002 ± 0.001 | 0.0008 – 0.22 | 0.02 ± 0.06 | 0.0009 – 0.03 | 0.01 ± 0.008 |
| P-a (mg.g^-1^) | 0.05 – 3.49 | 1.53 ± 1.56 | 0.06 – 3.69 | 0.90 ± 1.06 | 0.39 – 11.9 | 6.56 ± 4.93 |
| CEC (cmol_(+)_.kg^-1^) | 0.58 – 31.1 | 14.5 ± 13.0 | 2.02 – 15.7 | 6.58 ± 4.39 | 2.18 – 31.2 | 16.1 ± 10.5 |
| Ca^2+^ (cmol_(+)_.kg^-1^) | 0.34 – 24.3 | 11.3 ± 10.2 | 1.54 – 17.4 | 7.12 ± 4.98 | 1.76 – 28.1 | 15.1 ± 9.61 |
| Mg^2+^ (cmol_(+)_.kg^-1^) | 0.14 – 9.71 | 4.00 ± 3.52 | 0.62 – 3.18 | 1.57 ± 0.92 | 0.63 – 9.58 | 4.42 ± 3.21 |
| Na^+^ (cmol_(+)_.kg^-1^) | 0.03 – 0.95 | 0.36 ± 0.34 | 0.05 – 3.21 | 0.77 ± 1.02 | 0.05 – 4.85 | 1.67 ± 1.62 |
| K^+^ (cmol_(+)_.kg^-1^) | 0.007 – 0.29 | 0.10 ± 0.09 | 0.01 – 1.14 | 0.25 ± 0.36 | 0.02 – 1.46 | 0.43 ± 0.48 |

| Fixed effects | Mean square | df | F value | P value | Conditional R2 | Marginal R2 |
| --- | --- | --- | --- | --- | --- | --- |
| **% N** | | | | | | |
| Treatment | 2.341 | 2 | 23.276 | **0.041** | 0.70 | 0.68 |
| Season | 0.514 | 1 | 5.113 | **0.032** |  |  |
| Habitat | 0.320 | 1 | 3.180 | 0.217 |  |  |
| Treatment x Season | 0.147 | 2 | 1.457 | 0.251 |  |  |
| **% P** | | | | | | |
| Treatment | 0.0006 | 2 | 0.012 | 0.988 | 0.88 | 0.86 |
| Season | 0.166 | 1 | 33.420 | **< 0.0001** |  |  |
| Habitat | 0.364 | 1 | 73.296 | **0.014** |  |  |
| Treatment x Season | 0.002 | 2 | 0.431 | 0.654 |  |  |
| **log C:N** |  |  |  |  |  |  |
| Treatment | 0.163 | 2 | 8.468 | 0.106 | 0.77 | 0.64 |
| Season | 0.047 | 1 | 2.430 | 0.131 |  |  |
| Habitat | 0.141 | 1 | 7.307 | 0.114 |  |  |
| Treatment x Season | 0.052 | 2 | 2.713 | 0.084 |  |  |
| **log C:P** |  |  |  |  |  |  |
| Treatment | 0.005 | 2 | 0.137 | 0.880 | 0.91 | 0.86 |
| Season | 1.145 | 1 | 33.037 | **< 0.0001** |  |  |
| Habitat | 1.682 | 1 | 48.517 | **0.020** |  |  |
| Treatment x Season | 0.030 | 2 | 0.862 | 0.434 |  |  |
| **δ^13^C** |  |  |  |  |  |  |
| Treatment | 1.008 | 2 | 2.346 | 0.299 | 0.88 | 0.84 |
| Season | 72.024 | 1 | 167.595 | **< 0.0001** |  |  |
| Habitat | 7.255 | 1 | 16.883 | 0.054 |  |  |
| Treatment x Season | 5.566 | 2 | 12.952 | **< 0.0001** |  |  |

**Table S4**. Results of linear mixed models fitted to test changes in foliar nutrient parameters of coastal vegetation under different seabird influence (control, red footed booby, tern) and season (wet, dry), with habitat type (coastal shrub, grassland) as a covariate and sample location as a random effect. Statistically significant results (*p* ≤ 0.05) are indicated in bold. df: Degree of freedom.

**Table S5.** Results of linear mixed models fitted to test changes foliar nutrient parameters of seagrass under different seabird influence (control, red footed booby, tern) and season (wet, dry), with sample location as a random effect. Statistically significant results (*p* ≤ 0.05) are indicated in bold. df: Degree of freedom.

| Fixed effects | Mean square | df | F value | P value | Conditional R2 | Marginal R2 |
| --- | --- | --- | --- | --- | --- | --- |
| **% N** | | | | | | |
| Treatment | 0.092 | 2 | 10.155 | **0.002** | 0.82 | 0.71 |
| Season | 0.426 | 1 | 47.028 | **< 0.0001** |  |  |
| Treatment x Season | 0.212 | 2 | 23.385 | **< 0.0001** |  |  |
| **% P** | | | | | | |
| Treatment | 0.002 | 2 | 2.836 | 0.090 | 0.28 | 0.28 |
| Season | 0.002 | 1 | 3.573 | 0.078 |  |  |
| Treatment x Season | 0.001 | 2 | 2.189 | 0.146 |  |  |
| **log C:N** |  |  |  |  |  |  |
| Treatment | 0.052 | 2 | 7.750 | **0.005** | 0.71 | 0.58 |
| Season | 0.053 | 1 | 7.827 | **0.014** |  |  |
| Treatment x Season | 0.110 | 2 | 16.302 | **0.0002** |  |  |
| **log C:P** |  |  |  |  |  |  |
| Treatment | 0.064 | 2 | 3.843 | **0.045** | 0.30 | 0.30 |
| Season | 0.015 | 1 | 0.925 | 0.351 |  |  |
| Treatment x Season | 0.055 | 2 | 3.273 | 0.066 |  |  |
| **δ^13^C** |  |  |  |  |  |  |
| Treatment | 3.586 | 2 | 7.100 | **0.007** | 0.74 | 0.52 |
| Season | 12.876 | 1 | 25.492 | **0.0001** |  |  |
| Treatment x Season | 1.637 | 2 | 3.241 | 0.068 |  |  |
